# ParallelEvolCCM: Quantifying co-evolutionary patterns among genomic features

**DOI:** 10.1101/2024.06.12.598729

**Authors:** Robert G. Beiko, Chaoyue Liu, João Vitor Cavalcante, Ryan C. Fink

## Abstract

Concerted gains and losses of genomic features such as genes and mobile genetic elements can provide key clues into related functional roles and shared evolutionary trajectories. By capturing phylogenetic signals, a co-evolutionary model can outperform comparative methods based on shared presence and absence of features.We previously developed the Community Coevolution Model, which represents the gain/loss probability of each feature as a combination of its own intrinsic rate, combined the joint probabilities of gain and loss with all other features. Originally implemented as an R library, we have now developed a R wrapper that adds parallelization and several options to pre-filter the features to increase the efficiency of comparisons. Here we describe the functionality of EvolCCM and apply it to a dataset of 1000 genomes of the genus Bifidobacterium. ParallelEvolCCM is released under the MIT license and available at https://github.com/beiko-lab/arete/blob/master/bin/ParallelEvolCCM.R.

**Significance Statement:** Patchy phylogenetic distributions of genes, mobile genetic elements, and other genomic features can constitute evidence for lateral gene transfer. Comparing the presence/absence patterns of multiple features can reveal important associations among them, but the phylogenetic relationships must be taken into consideration in order to avoid spurious correlations. Our new ParallelEvolCCM software embeds these comparisons in a coevolutionary framework, offers a range of options to optimize the speed and comparisons, and offers helper scripts to visualize relationships among features.

## Introduction

The distribution of genomic features such as mutations, genes, and mobile genetic elements can be represented using presence/absence representations often referred to as phylogenetic profiles [Gaasterland and Ragan, 1998, Pellegrini et al., 1999]. Correlations and anti-correlations between these profiles can reveal a great deal about the capabilities, evolutionary history, and ecological roles of features and the implicated genomes. Although the term ‘phylogenetic’ is present in their name, these profiles do not account for the evolutionary relationships among their genomes. In many applications, this can lead to misleading patterns of correlation, especially in datasets with highly uneven sampling, such as those involving clinically important pathogens. Indeed, the unevenness of sampling across the bacterial tree is acute in references such as the database constructed from over 661,000 reference bacterial genomes, where the 20 most-abundant species (all with high representation of human pathogenic isolates) comprise over 90% of all sequenced genomes [Blackwell et al., 2021]. In these cases and in presence of high rates of lateral gene transfer among microorganisms, consideration of phylogenetic relationships can identify shared presence / absence patterns that are unexpected when compared to the reference tree of genomes. Several recent methods have aimed to correct for phylogenetic correlation in different ways including an approach based on Pagel’s co-evolutionary model [Liu et al., 2018], PhyloCorrelate [Tremblay et al., 2021], and a method that uses the inverse Potts model to minimize the impact of spurious transitive correlations among features [Fukunaga and Iwasaki, 2022].

Liu et al. [2023] introduced EvolCCM, an algorithm and R library that uses a co-evolutionary model to identify correlation patterns among features by modeling each feature’s rate of change in terms of its own intrinsic rate, combined with interactions with other features in the set. We demonstrated the accuracy of EvolCCM on simulated profiles with different degrees of association, and recovered key functional associations including shared GO terms and functional complexes of proteins. Although EvolCCM showed high levels of sensitivity, the inference of associations for all pairs of features is time consuming, scaling quadratically with the number of features and linearly with the size of the tree. Scaling EvolCCM to thousands of features and thousands of genomes will be infeasible; however, we can greatly expand its applicability by aggressively reducing inferred feature sets to those of greatest interest to the user. With this in mind, we have developed ParallelEvolCCM, an R wrapper that supports many types of feature filtering, uses parallelization to accelerate the generation of results, and provides a command-line interface that allows the user to invoke EvolCCM automatically. We demonstrate the utility of ParallelEvolCCM and its parameters by applying it to 1000 genomes from the genus *Bifidobacterium*, a group of Gram-positive, nonmotile bacteria with relatively small genomes and high GC content, which encompasses a diversity of species and strains that are associated with health and disease.

## Methods

ParallelEvolCCM uses the EvolCCM R library from Liu et al. [2023]. All phylogenetic operations are performed using the ape library [Paradis and Schliep, 2019]. EvolCCM models the transition rates of a set of binary features as a combination of their intrinsic rates and the association strength between features during the evolutionary process. Although the states in the original paper were based on the presence or absence of genes (i.e., phylogenetic profiles), EvolCCM can compare any set of features that have binary presence/absence states.

Given a vector *S*_*i*_ = *{x*_*i*,*k*_; *k* = 1, …, *n}* representing the system in state *i*, where *x*_*i*,*k*_ denotes the state of a feature *k* among *n* total features, the instantaneous transition rate *τ*_*i*,*k*_ for the specific gene *k* is defined as

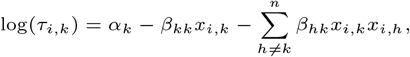

where *α*_*k*_ represents the intrinsic rate for feature *k*; *x*_*i*,*k*_ and *x*_*i*,*h*_ denote the current states of feature *k* and another feature *h* in the system, with values of –1 for absence and 1 for presence; *β*_*kk*_ indicates half the difference between the gain and loss rates of feature *k*; and *β*_*hk*_ is the coefficient of interaction between features *k* and *h*. The evolutionary changes in the system state along a phylogenetic tree are modeled as a continuous-time Markov process and the likelihood function *L* across the tree is constructed using Felsenstein’s pruning algorithm [Felsenstein, 1973]. The maximum likelihood estimates of the parameters in the transition rates are then obtained using the quasi-Newton method [Paradis and Schliep, 2019].

The estimated parameters provide insights into the evolutionary dynamics of the features. The intrinsic rate *α*_*k*_ represents the natural evolutionary base rate for feature *k*, and the interaction term *β*_*hk*_ demonstrates the strength and direction of the association between two features. To test the significance of these associations, a *Z*-test is performed to compute a p-value for the null hypothesis *H*_0_: *β*_*hk*_ = 0 (i.e., that the two features have no association with one another).

### Input

ParallelEvolCCM accepts as input a tab-separated feature file, where each row corresponds to a genome and each column a feature. ‘0’ in a given row/column combination indicates the absence of the corresponding feature from a genome, and ‘1’ its presence. The script also requires a Newick-formatted tree as input. EvolCCM requires a rooted tree: if the user-provided tree is unrooted, ParallelEvolCCM performs a midpoint rooting of the tree. All genome IDs specified in the feature file must have corresponding leaves in the tree; however not all leaf labels need to be present in the feature file. By default any multifurcations in the input tree are resolved randomly; a hardcoded option in the script can be changed to end execution if the provided tree is not binary.

### Parallelization

Parallelization is controlled through the ‘--cores’ option. Specifying a positive integer will attempt to use the corresponding number of CPU cores for the pairwise gene computations. Specifying ‘-1’ as a value will attempt to use all available cores on the system. If no value is specified, EvolCCM will use only a single core.

### Feature filtering

ParallelEvolCCM optionally uses two techniques to perform feature filtering. Features can be removed from consideration based on their abundance, with the logic that features that are found in all or nearly all genomes are unlikely to be interesting from a modeling point of view, and rare features will in most cases have minimal overlap in occurrence patterns, yielding no informative associations. Minimum and maximum abundance thresholds can be set independently, with any feature outside these thresholds removed from further consideration.

Subsets of features can be identified with common prefixes: for example, all plasmid-based annotations might be prefixed with “plasmid “, or all genes with a given Gene Ontology biological process ontology term with the ID of that term. Two flags, ‘--compare from’ and ‘--compare to’, determine which prefix-based subsets are to be compared. For example, ‘--compare from plasmid, genomicisland ‘ will compare only features with those prefixes against the full set of features. Similarly, ‘--compare from plasmid –-compare to plasmid’ will perform comparisons only within this group of features.

### Program output and visualization

ParallelEvolCCM provides detailed standard output about dataset dimensions, cores used, tree operations, and statistics about the number of filtered features. The tree used for rate inference after any midpoint rooting or resolution of multifurcating nodes is saved in an output file. Log-likelihoods and p-values for all pairwise feature comparisons are output as tab-separated matrices in two separate files. A script, ‘GraphML From EvolCCM.py’ can be used to generate GraphML files from either of the two matrices, for import into software packages such as Cytoscape [Shannon et al., 2003].

## Results

We retrieved 1000 assembled genomes assigned to the genus *Bifidobacterium* from the “AllTheBacteria” dataset [Hunt et al., 2024] that had a minimum N50 of 50,000 nucleotides and a length between 1.5 and 3.0 megabases, reflecting the approximate size bounds for the genus [Rodriguez and Martiny, 2020]. From this larger set we built a subset of 1000 genomes with species distributions (Table 1) that maintained some level of overrepresentation in the original dataset; for example, *Bifidobacterium longum* remained the most-common genome in the set. *Bifidobacterium* is often associated with positive gut health and dominates the gut flora of healthy, breast-fed infants (e.g., [Arboleya et al., 2016], but the genus also includes species such as *Bifidobacterium vaginale* (recently renamed from *Gardnerella vaginalis*, previously *Corynebacterium vaginale*, previously *Haemophilus vaginalis*: [Catlin, 1992]) that are associated with negative health outcomes such as bacterial vaginosis (e.g., [Morrill et al., 2020]). We also generated a subset of 100 genomes from the 1000-genome subset.

**Table 1.** Table 1. Distribution of most-common species in the 1000– and 100-genome *Bifidobacterium* datasets.

| Species name | B1000 | B100 |
| --- | --- | --- |
| *Bifidobacterium longum* | 300 | 25 |
| *Bifidobacterium adolescentis* | 150 | 22 |
| *Bifidobacterium pseudocatenulatum* | 100 | 7 |
| *Bifidobacterium breve* | 100 | 10 |
| *Bifidobacterium animalis* | 100 | 8 |
| *Bifidobacterium infantis* | 50 | 7 |
| *Bifidobacterium bifidum* | 50 | 5 |
| *Bifidobacterium vaginale* | 40 | 6 |
| Other | 110 | 10 |

We used the ARETE annotation and phylogenomics pipeline (https://github.com/beiko-lab/arete) to perform annotation and phylogenomic analysis of both these subsets independently. Distribution information was generated for several types of features: antimicrobial-resistance genes using version 6.0.2 of the Resistance Gene Identifier [Alcock et al., 2023], plasmids using MOB-suite [Robertson and Nash, 2018] version 3.0.3, and metal-resistance genes and virulence factors through homology search against the BacMet [Pal et al., 2014] and VFDB [Liu et al., 2021] databases using DIAMOND version 2.0.15 [Buchfink et al., 2021]. Reference phylogenetic trees were constructed from the concatenated core-genome sequence alignment generated by PPanGGOLiN v2.0.5 [Gautreau et al., 2020] using Fasttree version 2.1.10 [Price et al., 2010]. We then ran ParallelEvolCCM using various parameter settings to demonstrate runtimes and outputs. Comparisons between very rare features can sometimes fail to converge, leading to invalid small or large Chi-squared statistics; these were discarded prior to calculation of distributions.

Parallelization of comparisons into concurrent tasks showed the effectiveness of parallelization, as demonstrated in Figure 1. Each twofold increase in the number of cores yielded a speedup factor between 1.6 and 2.1, with the 100-genome run requiring between 5 minutes and 54 minutes, and the 1000-genome run requiring a minimum of 235 minutes and a maximum of 3072.

**Fig. 1.**
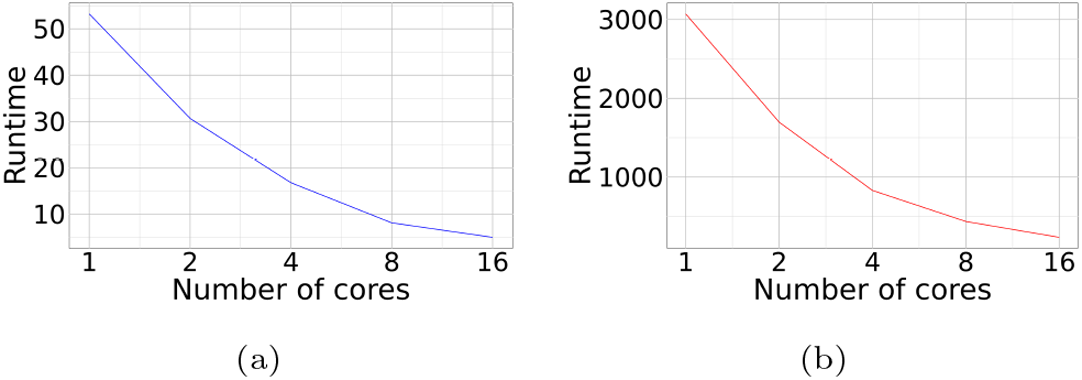
Runtimes in minutes for features extracted from 100 genomes (a) and 1000 genomes (b) with different numbers of allocated cores.

Figure 2 shows the feature and statistical distributions for the 100-genome and 1000-genome datasets. A total of 138 features (13 plasmids, 115 AMR genes, 2 BacMet, 8 VFDB) were observed at least once in the 100-genome dataset (Figure 2a). The 1000-genome dataset had a total of 384 predicted features (40 plasmids, 317 AMR genes, 8 BacMet, 19 VFDB); 333 of these were present in fewer than 50 genomes and 256 in fewer than five (Figure 2e). Chi-squared interaction scores in the 100-genome dataset ranged from –2.58 to 6.43, and the smallest p-value of 1.26*×*10^−10^ was obtained from a comparison between the ‘OCH.4’ and ‘TEM.116’ profiles (Figure 2b-d). The 1000-genome dataset had a larger range of –7.22 to 8.01, although the standard deviations of scores were similar between the two datasets (0.71 vs 0.76). The smallest p-value was 1.11*×*10^−15^, obtained from a comparison of plasmid AF432 and the IND.2a AMR gene. Restricting the 1000-genome dataset analysis to features with abundance between 5% and 95% required less than ten minutes when 16 CPU cores were used (Figure 2f-h).

**Fig. 2.**
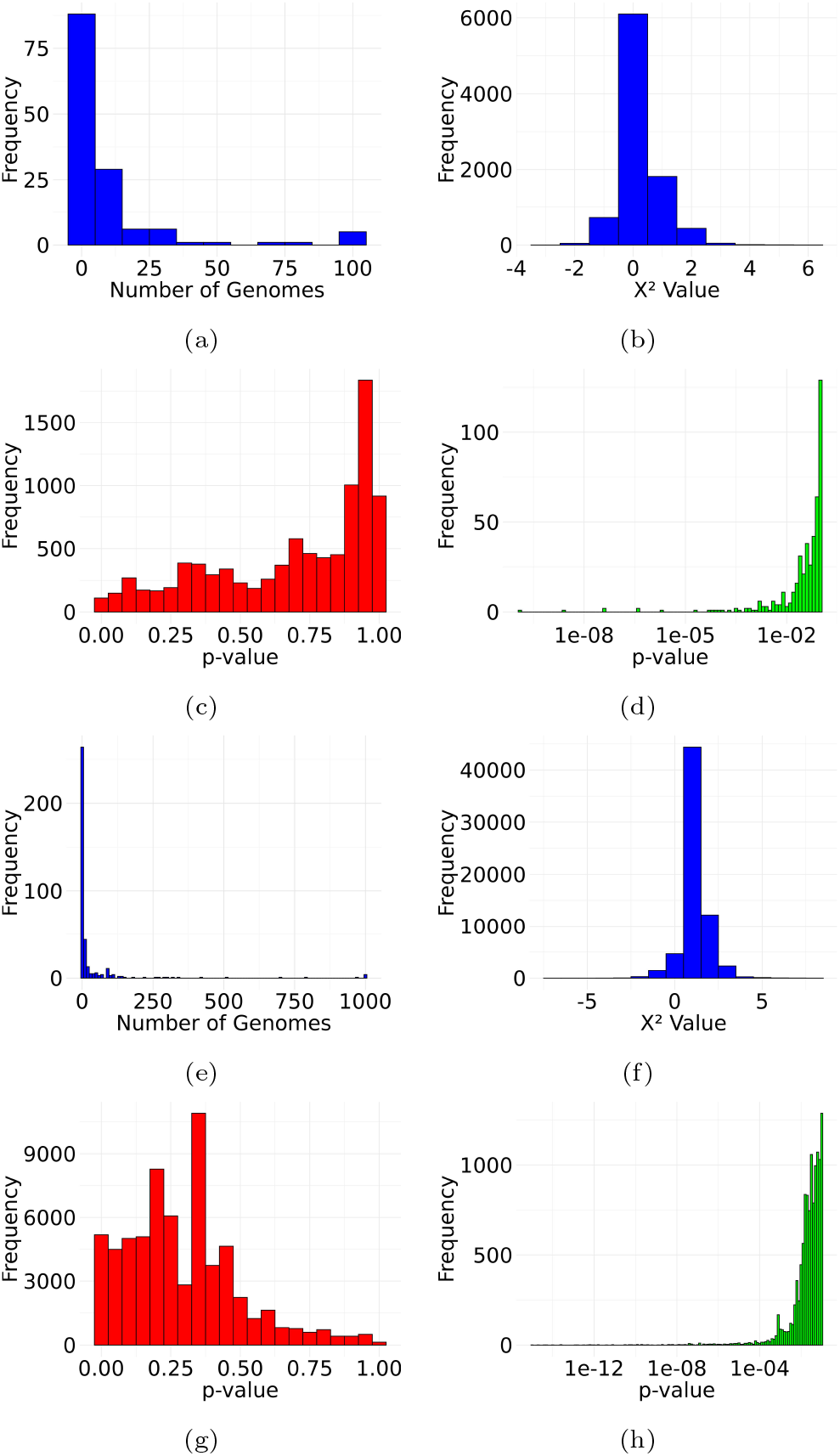
Feature and statistical score distributions for 100-genome (a-d) and 1000-genome (e-h) datasets. (a/e) Frequency distribution of feature counts across genomes. (b/f) Distribution of Chi-squared scores across all pairwise feature comparisons. (c/g) p-value distribution across pairwise comparisons on a linear scale from p = 0 to 1. (d/h) distribution of p-values less than or equal to 0.1 on a logarithimic scale.

Figure 3 shows thresholded EvolCCM networks visualized in Cytoscape. Thresholding the features in the 100-genome data set at a p-value of 0.005 (Figure 3a) yielded a total of three connected components comprising two features (*ureB* and *ureG*), three features (*nimG, APH*.*4*..*1a*, and *ranB*), and 28 features. *ugd* is present in both VFDB and CARD, and the two features were connected to one another. A total of five MOB-suite plasmid clusters showed strong associations with other features; one set of plasmids showed connections with the beta-lactamases SPN79.1 and*cfxA*, and the *tap* efflux pump, while the other showed connections to the beta-lactamases OXA.118, OXA.569, and OCH.4, as well as the colistin-resistance gene MCR 7.1.

**Fig. 3.**
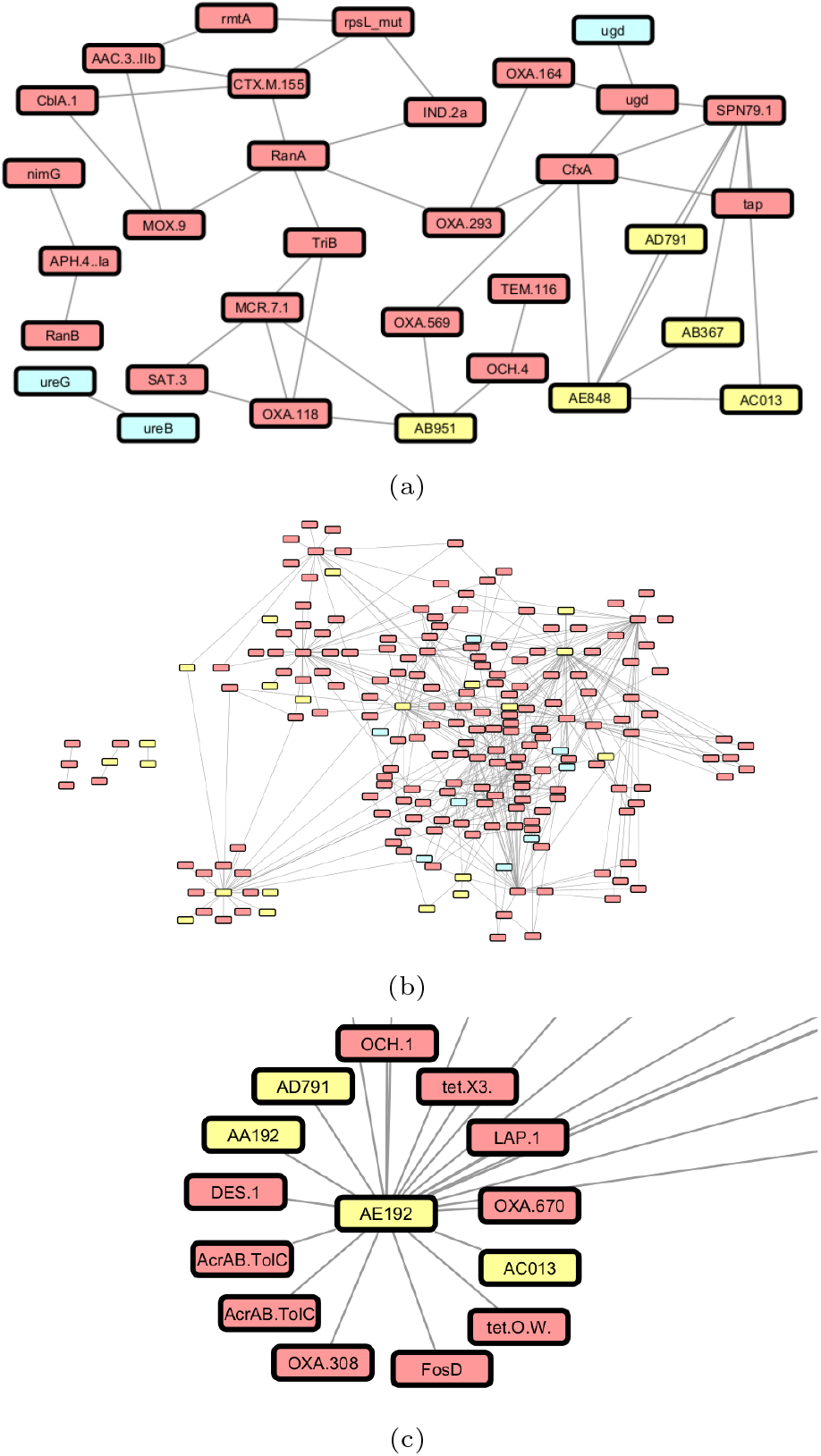
Network visualizations of associations inferred using EvolCCM. Nodes represent annotated plasmids (yellow), AMR genes (red), and VFDB/BacMet genes annotated through homology search (cyan). (a) Connections between features in the 100-genome dataset with associated p-values less than 0.005. (b) Connections between features in the 1000-genome dataset with associated p-values less than 0.005. (c) Subgraph from (b), showing nodes connected to plasmid AE192.

Nearly all features in the 1000-genome dataset were in a single connected component, with three additional components comprising three, three, and two features respectively (Figure 3b). Plasmid cluster AE192 was a hub in the large connected component, and was the only feature to which 13 others were connected. Three of these were plasmids, with the remaining ten comprising beta-lactamases (OCH.1, OXA.670, OXA.308, and DES.1), the cephalosporin-resistance-conferring LAP.1, the tetracycline-resistance genes tet.X3 and tet.O.W, fosfomycin resistance *fosD*, and the multidrug efflux pump AcrAB.TolC (Figure 3c). Cluster AE192 has a single reference plasmid with GenBank accession AB646744.1 from *Bacteroides fragilis* with size 12,817 base pairs, which contains multiple resistance proteins [Goto et al., 2012]. Although this reference plasmid is too small to contain all the associated genes in the implied EvolCCM network, the plasmid predicted to occur in *Bifidobacterium* may have different properties which could be investigated by inspecting the sequence assemblies in this dataset.

## Conclusions

The coevolutionary model of EvolCCM performs comparisons of feature profiles with appropriate corrections for phylogenetic correlations. The need for this correction is highest when dataset sampling is uneven, which is frequently the case in intensively sampled species that may contain one or more sets of highly similar outbreak isolates. ParallelEvolCCM can accept tables of features generated from any source and offers a range of options to accelerate computations through parallelization and reduction in the number of pairwise comparisons that need to be done. We previously demonstrated that the “Community” aspect of EvolCCM, which allows comparisons of sets of more than two features at a time, can effectively filter out false-positive connections. Considering sets of size *>*2 increases the complexity and number of comparisons, and ParallelEvolCCM offers an opportunity to accelerate these investigations as well.

## Supporting information

Documentation, source code, and example files used in the manuscript.

## Acknowledgments

This work was supported by Genome Canada, Research Nova Scotia, the Natural Sciences and Engineering Research Council of Canada, the Dalhousie Faculty of Computer Science, and Genome Atlantic, with computing support from the Digital Research Alliance of Canada.

## Author contributions statement

R.G.B. wrote the ParallelEvolCCM script, performed testing, and generated the results. C.L. developed the EvolCCM model and performed refinements to the latest version. All authors conceived the experiments, provided conceptual input, and contributed to the paper. A first draft of some parts of the EvolCCM code was produced using ChatGPT, with extensive subsequent modifications and testing by R.G.B.

## Software and Data Availability

EvolCCM and ParallelEvolCCM are released under the MIT license. The source code for ParallelEvolCCM is available at https://github.com/beiko-lab/arete/blob/master/bin/ParallelEvolCCM.R. Documentation for ParallelEvolCCM can be found at https://beiko-lab.github.io/arete/evolccm/. An archive containing the input files, documentation, ParallelEvolCCM results, and helper scripts used to generate Figures 2 and 3 is available at the same location as the R script. The feature profiles include the accession numbers for all genomes used in this study.

