## Supplementary material for "ParallelEvolCCM: Quantifying co-evolutionary patterns among genomic features": Documentation, source code, and example files used in the manuscript.: ParallelEvolCCM_Doc.pdf

### ParallelEvolCCM Release Package

Rob Beiko

June 10, 2024

“ParallelEvolCCM” is abbreviated as “PECCM” in this document because it’s faster to type.

#### 1 Package Contents

The package contains three subfolders:

**Scripts/** The R scripts used to generate feature and statistical histograms, and the Python script that is used to build the GraphML files from PECCM output.

**100Bifido/** The results of the 100-genome analysis described in the paper.

**1000Bifido/** The results of the 1000-genome analysis described in the paper.

Each of the results folders contains the following subdirectories and files. X is a placeholder for the size (i.e., 100 or 1000):

**SourceFiles/** The source files.

- Bifido_X.tre: The Newick-formatted tree
- Bifido_X_feature_profile: The tab-separated feature file. **This is the input file for ‘PECCM_BuildFeatureHistogram.R’, see usage below**

**Results/** The results produced by PECCM and the helper scripts.

- EvolCCM_Bifido_X.tre: The tree used by EvolCCM (with midpoint rooting and multifurcating node resolution if necessary)
- EvolCCM_Bifido_X_feature_profile.tsv: A tab-separated file with the p-values and statistics for all pairwise comparisons between features. **This is the input file for the scripts ‘PECCM_BuildStatHistogram.R’ and ‘PECCM_Build_GraphML.py’, see below**
- EvolCCM_Bifido_X_feature_profile.tsv.pvals and EvolCCM_Bifido_X_feature_profile.tsv.pvals: Tab-separated matrices showing the p-values and $X^2$ scores for all features.
- EvolCCM_Bifido_100.graphml: GraphML-formatted file with connections between features.
- Four .jpg files: ‘a_’ is the output feature profile, and ‘b_’, ‘c_’, and ‘d_’ are the feature and statistical distributions.

### 2 Usage

The examples below are based on the '100Bifido' dataset; it suffices to change to the other directory and add another zero if you would like to work with the 1000-genome dataset.

#### 2.1 Running ParallelEvolCCM

ParallelEvolCCM.R has several dependencies, which should automatically be installed the first time you run the script. You may need to install missing Linux packages using the following command:

```
sudo apt-get install libssl-dev libfontconfig1-dev libharfbuzz-dev
libfribidi-dev libfreetype6-dev libpng-dev libtiff5-dev libjpeg-dev
libopenblas-dev
```

Here are the options for the script:

```
RunEvolCCM.R --intree <tree_file> --intable <table_file> [--compare_from
value1,value2,...]  [--compare_to valueA,valueB,...]  [--cores
<number_or_-1>] [--min_abundance <0.0-1.0>] [--max_abundance <0.0-1.0>]
```

Options:<br>
--intree <tree_file> : Path to the tree file (required).<br>
--intable <table_file> : Path to the table file (required).<br>
--compare_from <values> : Comma-separated list of values for comparison (optional).<br>
--compare_to <values> : Comma-separated list of values for comparison (optional).<br>
--cores <number_or_-1> : Specify number of cores for parallel processing or '-1' for all cores (optional, default is 1).<br>
--min_abundance <0.0-1.0> : Minimum abundance proportion threshold (optional, default is 0.0).<br>
--max_abundance <0.0-1.0> : Maximum abundance proportion threshold (optional, default is 1.0).<br>
--show_nans : Show NaNs in matrix (default is to convert non-converging $X^2$ values to 0 and p-values to 1.<br>
-h : Show this help message.

This command (specifying any reasonable number of cores) will recreate the results of the 100-genome dataset:

```
Rscript ParallelEvolCCM.R --intree Bifido_100.tre --intable
Bifido_100_feature_profile.tsv --min_abundance 0.05 --max_abundance 0.95
--cores 8
```

Four output files will be produced. All will be prefixed with 'EvolCCM_' to distiguish them from the input files.

- .tre file: The tree used by EvolCCM (with midpoint rooting and multifurcating node resolution if necessary). This file will end with a '.tre' extension.
- .tsv file: Statistics associated with the EvolCCM comparisons, with one line for each pairwise comparison.
- .tsv.pvals file: A matrix showing the p-values from all-versus-all comparisons between features.
- .tsv.X2 file: A matrix showing the $X^2$ values from all-versus-all comparisons between features.

#### 2.2 Helper scripts

**PECCM_BuildFeatureHistogram.R** can be used to generate the feature distribution histogram. Usage is:

```
Rscript PECCM_BuildFeatureHistogram.R infile
```

Where 'infile' is the input feature table (for example, Bifido_100_feature_profile.tsv). There are no other command-line options.

A single .jpg file will be produced.

**PECCM_BuildStatHistogram.R** is used to generate the statistical summary histograms. Usage is:

```
Rscript PECCM_BuildStatHistogram.R infile
```

Where 'infile' is the input table of results ('EvolCCM...tsv').
Three .jpg files will be produced.

**PECCM_Build_GraphML.py** is used to generate a graph from the pairwise comparisons, with optional p-value thresholding. You can also use the –attribute_name_length option to truncate attribute names for visual purposes.

The optional 'type_underscore' argument will treat the first part of each attribute name (up to the first underscore) as its type: for example, 'plasmid_ABC' and 'plasmid_def' would both be treated as objects of type 'plasmid', with names 'ABC' and 'DEF', respectively.

```
Usage:  PECCM_Build_GraphML.py [-h] [--p_value_threshold P_VALUE_THRESHOLD]
[--attribute_name_length ATTRIBUTE_NAME_LENGTH] [--type_underscore]
input_file output_file
```

Here is an example to generate the 100-genome graph:

```
python ../PECCM_Build_GraphML.py --attribute_name_length
10 --type_underscore EvolCCM_Bifido_100_feature_profile.tsv
EvolCCM_Bifido_100.graphml
```
