## Supplementary material for "ParallelEvolCCM: Quantifying co-evolutionary patterns among genomic features": Documentation, source code, and example files used in the manuscript.: ParallelEvolCCM.R

```
### ParallelEvolCCM.R
### Hosted at: https://github.com/beiko-lab/arete/blob/master/bin/ParallelEvolCCM.R
### Version 1.0, released *** June 2024
### Released under MIT License (https://opensource.org/license/mit)
### Note: igraph requires the OpenBlas library, which can be installed with:
### sudo apt-get install libopenblas-dev
### devtools: sudo apt-get install libssl-dev libfontconfig1-dev libharfbuzz-dev libfribidi-dev libfreetype6-dev libpng-dev libtiff5-dev libjpeg-dev

# List of CRAN packages to check and install
cran_packages <- c("ape", "dplyr", "devtools", "phytools", "foreach", "doParallel", "gplots")

for (package in cran_packages) {
    if (!suppressPackageStartupMessages(require(package, character.only = TRUE))) {
        install.packages(package)
    }
	suppressPackageStartupMessages(library(package, character.only = TRUE))
}

if (!require(phytools)) {
	remotes::install_github("liamrevell/phytools")
}

if (!require(evolCCM)) {
	devtools::install_github("beiko-lab/evolCCM")
}

suppressPackageStartupMessages(library(evolCCM))


#####################################################################
### FUNCTION PARSE_ARGS
### Parse the command-line arguments.
### args: the arguments
### RETURN a list that contains values for all the defined arguments.
#####################################################################

parse_args <- function(args) {

  # Check if no arguments are provided or if '-h' flag is given
  if (length(args) == 0 || any(args == "-h")) {
    cat("Usage: RunEvolCCM.R --intree <tree_file> --intable <table_file> [--compare_from value1,value2,...] [--compare_to valueA,valueB,...] [--cores <number_or_-1>] [--min_abundance <0.0-1.0>] [--max_abundance <0.0-1.0>]\n")
    cat("Options:\n")
    cat("  --intree <tree_file>        : Path to the tree file (required).\n")
    cat("  --intable <table_file>      : Path to the table file (required).\n")
    cat("  --compare_from <values>     : Comma-separated list of values for comparison (optional).\n")
    cat("  --compare_to <values>       : Comma-separated list of values for comparison (optional).\n")
    cat("  --cores <number_or_-1>      : Specify number of cores for parallel processing or '-1' for all cores (optional, default is 1).\n")
    cat("  --min_abundance <0.0-1.0>   : Minimum abundance proportion threshold (optional, default is 0.0).\n")
    cat("  --max_abundance <0.0-1.0>   : Maximum abundance proportion threshold (optional, default is 1.0).\n")
	cat("  --show_nans                 : Show NaNs in matrix (default is to convert non-converging X^2 values to 0 and p-values to 1.\n")
    cat("  -h                          : Show this help message.\n")
    quit(save="no")
  }

  cat("\n\n*********************** PARALLELEVOLCCM ***********************\n\n")
  
  cat("Parsing arguments...\n")
  
  # Function to parse command line arguments without '='
  parseArg <- function(argname, options) {
    index <- which(options == argname)
    if (length(index) > 0 && index < length(options)) {
      return(options[index + 1])
    }
    NULL
  }
  
  # Parse the CL arguments
  inputTree <- parseArg("--intree", args)
  inputProfile <- parseArg("--intable", args)
  compare_to_str <- parseArg("--compare_to", args)
  min_abundance_str <- parseArg("--min_abundance", args)
  max_abundance_str <- parseArg("--max_abundance", args)
  cores_str <- parseArg("--cores", args)
  show_nans <- "--show_nans" %in% args
  
  # Parse the optional --compare_from and --compare_to arguments
  compare_from_str <- parseArg("--compare_from", args)
  if (!is.null(compare_from_str)) {
    compare_from_vector <- strsplit(compare_from_str, ",")[[1]]
  } else {
    compare_from_vector <- NULL
  }
  
  if (!is.null(compare_to_str)) {
    compare_to_vector <- strsplit(compare_to_str, ",")[[1]]
  } else {
    compare_to_vector <- NULL
  }

  # Set default values
  min_abund <- 0.0
  max_abund <- 1.0
  
  # Validate and assign min_abundance if specified
  if (!is.null(min_abundance_str)) {
    temp_val <- as.numeric(min_abundance_str)
    if (!is.na(temp_val) && temp_val >= 0.0 && temp_val <= 1.0) {
      min_abund <- temp_val
    } else {
      stop("--min_abundance must be a proportional value between 0.0 and 1.0", call. = FALSE)
    }
  }
  
  # Validate and assign max_abundance if specified
  if (!is.null(max_abundance_str)) {
    temp_val <- as.numeric(max_abundance_str)
    if (!is.na(temp_val) && temp_val >= 0.0 && temp_val <= 1.0) {
      max_abund <- temp_val
    } else {
      stop("--max_abundance must be a proportional value between 0.0 and 1.0", call. = FALSE)
    }
  }
  
  # Check the relationship between min_abund and max_abund
  if (min_abund > max_abund) {
    stop("--min_abundance value must be less than or equal to --max_abundance value", call. = FALSE)
  }
  
  # Determine the parallel processing parameters
  if (is.null(cores_str)) {
    num_cores <- 1
  } else if (cores_str == "-1") {
    num_cores <- detectCores()
  } else {
    num_cores <- as.integer(cores_str)
    if (is.na(num_cores) || num_cores <= 0) {
      stop("--cores argument must be a positive integer or '-1' for all cores", call. = FALSE)
    }
  }
  
  # Check if both --intree and --intable are provided
  if (is.null(inputTree) || is.null(inputProfile)) {
    stop("Usage: RunEvolCCM.R --intree <tree_file> --intable <table_file> [--compare_from value1,value2,...] [--compare_to valueA,valueB,...]", call. = FALSE)
  }

  parsed_list <- list("inputTree" = inputTree,
                      "inputProfile" = inputProfile,
                      "compare_from_vector" = compare_from_vector,
                      "compare_to_vector" = compare_to_vector,
                      "min_abund" = min_abund,
                      "max_abund" = max_abund,
					  "show_nans" = show_nans,
					  "num_cores" = num_cores)
  
  return(parsed_list)
}

##########################################################################
### FUNCTION MODIFY_TREE
### Apply fixes, if necessary, to make the tree suitable for CCM analysis.
### tree: the tree
### fix_multifurcations, coerce_branches: self-explanatory Booleans
### RETURNS the corrected tree
##########################################################################

modify_tree <- function(tree,fix_multifurcations,coerce_branches) {

  if (!is.rooted(tree)) {
    cat(noquote("*** Input tree is not rooted! Applying midpoint rooting algorithm. ***\n"))
    tree <- midpoint.root(tree)
  }
  
  ### Arbitrarily resolve multifurcations and zero-length branches ###
  if (!is.binary(tree)) {
    if (fix_multifurcations) {
      cat(noquote("*** Input tree is not binary! Forcing the issue. ***\n"))
      tree<-multi2di(tree)
    } else {
      stop("Tree contains multifurcations and \"fix_multifurcations\" is set to FALSE. Exiting...")
    }
  }
  
  ### Coerce branches if necessary ###
  if (coerce_branches) {
    short_branch_count <- sum(tree$edge.length < min_branch_length)
    if (short_branch_count > 0) {
      notification_string = paste(short_branch_count, " of ",nrow(tree$edge)," edges are less than the specified threshold of ",min_branch_length,".", sep = "")
      warning(paste(notification_string,"Setting these branches to equal the threshold; this will affect the results!"))
      tree$edge.length[tree$edge.length < min_branch_length] <- min_branch_length
    }
  }
  return(tree)
}

########################################################################################
### FUNCTION CHECK_AND_FILTER_PROFILE
### Check for consistency between the profile and the tree, and apply filtering criteria
### aprofile: the phylogenetic profile
### tree: the phylogenetic tree
### min_abund: minimum abundance threshold
### max_abund: maximum abundance threshold
### RETURN a filtered profile, as long as nothing is broken
########################################################################################

check_and_filter_profile <- function(aprofile,tree,min_abund,max_abund) {
  
  # Get the leaf labels from the phylogenetic tree
  tree_labels <- tree$tip.label
  
  # Get the row labels from the dataframe
  df_labels <- rownames(aprofile)
  
  mismatch <- 0
  # Find labels that are in the tree but not in the dataframe
  missing_in_df <- setdiff(tree_labels, df_labels)
  if (length(missing_in_df) > 0) {
    cat("Labels in the tree but not in the dataframe:",quote=FALSE)
    cat(missing_in_df)
    mismatch = 2
  }
  
  # Find labels that are in the dataframe but not in the tree
  missing_in_tree <- setdiff(df_labels, tree_labels)
  if (length(missing_in_tree) > 0) {
    cat("Labels in the dataframe but not in the tree:",quote=FALSE)
    cat(missing_in_tree)
    mismatch = 1
  }
  
  if (mismatch == 1) {
    stop("Exiting...")
  }
  
  cat("All names match between tree and matrix.\n")
  
  ### ABUNDANCE FILTER ###
  
  aprofile <- aprofile[tree$tip.label,]
  aprofile[aprofile > 1] <- 1
  
  numFeatures <- dim(aprofile)[2]
  cat(paste("Total number of features: ",numFeatures, "\n"))
  
  ### Abundance filter: subset the dataframe to include only those columns that satisfy the proportion criteria ###
  proportion_ones <- apply(aprofile, 2, function(col) sum(col == 1) / length(col))
  aprofile <- aprofile[, proportion_ones >= min_abund & proportion_ones <= max_abund]
  numFeatures <- dim(aprofile)[2]
  cat(paste("Total number of features post abundance filtering: ",numFeatures,"\n"))

  ### CLADE FILTERING: Remove any feature that maps perfectly onto a clade in the rooted phylogenetic tree ###
  # This function will recursively explore the tree and return the clades
  # Function to get all descendant tips of a node
  #if(clade_filter) {
  if(FALSE) {
    get_descendant_tips <- function(tree, node) {
      if (node <= length(tree$tip.label)) {
        return(tree$tip.label[node])
      } else {
        descend <- which(tree$edge[, 1] == node)
        tips <- c()
        for (i in descend) {
          tips <- c(tips, get_descendant_tips(tree, tree$edge[i, 2]))
        }
        return(tips)
      }
    }
    
    # Get the clades for all internal nodes
    clades <- lapply((length(tree$tip.label) + 1):(length(tree$tip.label) + tree$Nnode), 
                     function(node) get_descendant_tips(tree, node))
    
    # Function to check if a feature is unique to a clade
    check_feature <- function(feature){
      for(clade in clades){
        clade_data <- aprofile[unlist(clade), feature]
        outside_clade_data <- aprofile[setdiff(rownames(aprofile), unlist(clade)), feature]
        
        if (all(clade_data == 1, na.rm = TRUE) & all(outside_clade_data == 0, na.rm = TRUE)){
          return(TRUE)
        }
      }
      return(FALSE)
    }
    
    for(feature in colnames(aprofile)){
      if(check_feature(feature)){
        filtered_data <- filtered_data %>% select(-feature)
      }
    }
  }
  return(aprofile)
}

####################################################################
### FUNCTION PROCESS_PAIR
### Compute EvolCCM distance for a pair of profiles
### i,j: indices
### filtered_data: dataframe with phylogenetic profiles
### tree: the phylogenetic tree
### RETURNS a row containing CCM statistics
####################################################################

process_pair <- function(i, j, filtered_data, tree) {
  res <- tryCatch({
    #if (
    if (j %% 50 == 0) {
      print(paste(i,"vs",j))
    }
    
    # Run EstimateCCM for the current pair
    aE <- EstimateCCM(filtered_data[, c(i, j)], phytree = tree, trace = FALSE)
    estimatedRates <- aE$nlm.par
    
    # Retrieve the score and pvalue
    paE <- ProcessAE(aE)
    
    interact_score <- estimatedRates[5] / paE$hessianSE[5]
    interact_pval <- 2 * (1 - pnorm(abs(interact_score)))
    
    res <- c(aE$nlm.par, interact_score, interact_pval)
    
    # Return the result along with i, j, and column names
    list(i = i, j = j, colnames = colnames(filtered_data[, c(i, j)]), res = res)
    #}
  }, error = function(e) {
    # Return NULL in case of an error
    NULL
  })
  
  return(res)
}

###########################################################################
### FUNCTION CONSTRUCTMATRIX
### Build square matrices that represent similarities/distances of profiles
### lines: the output lines generated by CCM
### outputFile: the output file
### column: the column in 'lines' to use for matrix construction
### RETURNS nothing, WRITES the resulting matrix to a file
##########################################################################

constructMatrix <- function(lines, outputFile, column) {
  
# Initialize an empty list to store the entities
  entities <- list()
  
  for (line in lines) {
    # Parse the line into fields
    fields <- strsplit(line, "\t")[[1]]
    # Extract the entities
    entity1 <- fields[1]
    entity2 <- fields[2]

    # If the entities are new, add them to the list
    if (!entity1 %in% entities) {
      entities <- c(entities, entity1)
    }
    if (!entity2 %in% entities) {
      entities <- c(entities, entity2)
    }
  }
  
  # Initialize an empty matrix to store the distances
  distances <- matrix(nrow = length(entities), ncol = length(entities))
  
  for (line in lines) {
    # Parse the line into fields
    fields <- strsplit(line, "\t")[[1]]
    # Extract the entities and the distance
    entity1 <- fields[1]
    entity2 <- fields[2]
    distance <- as.numeric(fields[column])
    
    # Add the distance to the matrix
    distances[entities == entity1, entities == entity2] <- distance
    distances[entities == entity2, entities == entity1] <- distance
  }
  
  # Convert the matrix to a data frame and set the row and column names
  distances <- as.data.frame(distances)
  rownames(distances) <- entities
  colnames(distances) <- entities
  
  # Write the matrix to a file
  write.table(distances, file = outputFile, sep = "\t", quote = FALSE)
  
}

####################
### MAIN PROGRAM ###
####################


########################################## FILTERS ##########################################

### Remove any profile that maps perfectly onto a clade in the tree. In practice this doesn't do much ###
clade_filter = FALSE

### Apply pairwise heuristic based on some criterion: do not perform full comparison if heuristic doesn't satisfy the threshold ### 
### (CURRENTLY NOT IMPLEMENTED: mutual information and Euclidean were bad, UniFrac might be worth a try) ###
heuristic_filter = FALSE
heuristic_threshold = 0.001

### Coerce branches to a minimum threshold length (ALTERNATIVE: multiply all branch lengths by the minimum amount needed to get to threshold)
coerce_branches = FALSE
#min_branch_length = 0.001

### Coerce multifurcations to bifurcations (alternative is to stop execution)
fix_multifurcations = TRUE

######################################## PARSE CL ARGUMENTS ########################################

args <- commandArgs(trailingOnly = TRUE)
parsed <- parse_args(args)

### Output file name (could be changed to a command-line argument) ###
outputTree <- paste("EvolCCM_", basename(parsed$inputTree), sep = "")
outputFile <- paste("EvolCCM_", basename(parsed$inputProfile), sep = "")

###################### READ THE TREE FILE AND FIX IF NECESSARY ######################

cat("\nReading tree...\n")
tree <- read.tree(parsed$inputTree)
tree <- modify_tree(tree,fix_multifurcations,coerce_branches)
write.tree(tree,outputTree)

###################### READ THE PROFILE AND APPLY FILTERS ######################

### Read the profile file ###

cat("\nReading profile...\n")
aprofile <- read.csv(parsed$inputProfile, row.names=1,sep="\t")
filtered_data <- check_and_filter_profile(aprofile, tree, parsed$min_abund, parsed$max_abund)
numFeatures <- dim(filtered_data)[2]

######################## PARALLELIZED NESTED LOOP FOR PAIRWISE PROFILE COMPARISONS ########################

cat("\nInitiating comparisons...\n")
registerDoParallel(cores=parsed$num_cores)

compare_from_vector <- parsed$compare_from_vector
compare_to_vector <- parsed$compare_to_vector

results <- foreach(i = 1:(numFeatures - 1), .combine = 'c', .multicombine = TRUE) %dopar% { 
  i_name <- colnames(filtered_data)[i]
    
  # If compare_from_str is defined, then check the condition. If not, then set to TRUE by default.
  i_condition <- ifelse(!is.null(compare_from_vector),
                        any(sapply(compare_from_vector, function(x) {startsWith(i_name, x)})),
                        TRUE)
    
  if (i_condition) {
    print(paste("Processing feature #", i, " (", i_name, ")", sep = ""), quote = FALSE)
    
    sapply((i + 1):numFeatures, function(j) {
      j_name <- colnames(filtered_data)[j]
      
      # If compare_to_str is defined, then check the condition. If not, then set to TRUE by default.
      j_condition <- ifelse(!is.null(compare_to_vector),
                            any(sapply(compare_to_vector, function(x) {startsWith(j_name, x)})),
                            TRUE)
      
      if (j_condition) {
        return(process_pair(i, j, filtered_data, tree))
      } else {
        return(NULL)
      }
    }, simplify = FALSE)
  } else {
    return(NULL)
  }
}

cat("\nProfile comparisons complete!\n")

################################# OUTPUT #####################################

cat("\nWriting output...\n")

### Process the results and write them to chunks of output files ###
chunk_size <- 100
output_file_pattern <- "output_temp_chunk_%d.txt"
chunk_index <- 1
result_counter <- 1

for (result in results) {
  if (!is.null(result)) {
    if ((result_counter - 1) %% chunk_size == 0) {
      temp_output_file <- sprintf(output_file_pattern, chunk_index)
      chunk_index <- chunk_index + 1
    }
	### Fix NaNs unless user has specified otherwise
	if (parsed$show_nans != 1) {
		if (is.nan(result$res[6])) {
			# print(result$res)
			# print("Fixing NaN")
			result$res[6] = 0
			result$res[7] = 1
			# print(result$res)
		}
	}
    cat(paste(result$colnames,collapse="\t"), file = temp_output_file, append = TRUE)
    cat("\t", file = temp_output_file, append = TRUE)
    cat(paste(result$res,collapse="\t"), file = temp_output_file, append = TRUE)
    cat("\n", file = temp_output_file, append = TRUE)

    result_counter <- result_counter + 1
  }
}

# Merge the chunks of output files into the main output file
temp_files <- list.files(pattern = "output_temp_chunk_[0-9]+\\.txt")

outputHeadings <- paste("feature1","feature2","intrisic1", "intrisic2", "gainloss1", "gainloss2", "interaction", "interact_score", "interact_pval",sep="\t")
cat(outputHeadings, file = outputFile, sep = "\n")
for (temp_file in temp_files) {
  # Read the contents of the temporary file
  temp_contents <- readLines(temp_file)

  # Write the contents to the main output file
  cat(temp_contents, file = outputFile, append = TRUE, sep = "\n")

  # Remove the temporary file
  file.remove(temp_file)
}

# Read the lines from the input file
lines <- readLines(outputFile)[-1]
X2file = paste(outputFile,"X2",sep = ".")
Pfile = paste(outputFile,"pvals",sep = ".")

constructMatrix(lines,X2file,8)
constructMatrix(lines,Pfile,9)

cat("ParallelEvolCCM run done.\n\n")
```
