## Supplementary material for "ParallelEvolCCM: Quantifying co-evolutionary patterns among genomic features": Documentation, source code, and example files used in the manuscript.: PECCM_Build_GraphML.py

```
import argparse
import re
from xml.etree.ElementTree import Element, SubElement, tostring, ElementTree as ET
from pygraphml import Graph
from pygraphml import GraphMLParser

def shorten_attribute_name(attribute_name, length):
    return attribute_name[:length]

def escape_special_characters(attribute_name):
    return re.sub(r'[&<>\'"]', lambda c: f'&#{ord(c.group(0))};', attribute_name)

### Create nodes in the graph ###
def create_node(node_type, attribute_name, shortened_attribute_name):
    node = Element('node')
    node.set('type', node_type)
    
    full_attribute = SubElement(node, 'data')
    full_attribute.set('key', attribute_name)
    full_attribute.text = str(attribute_name) if attribute_name is not None else ''
    
    short_attribute = SubElement(node, 'data')
    short_attribute.set('key', shortened_attribute_name)
    short_attribute.text = str(shortened_attribute_name) if shortened_attribute_name is not None else ''
    
    return node

### Create edges in the graph ###
def create_edge(source, target, edge_type, x2_value, magnitude, sign, p_value):
    edge = Element('edge')
    edge.set('source', source)
    edge.set('target', target)
    
    edge_type_attr = SubElement(edge, 'data')
    edge_type_attr.set('key', 'edge_type')
    edge_type_attr.text = str(edge_type) if edge_type is not None else ''
    
    x2_value_attr = SubElement(edge, 'data')
    x2_value_attr.set('key', 'x2_value')
    x2_value_attr.text = str(x2_value) if x2_value is not None else ''
    
    magnitude_attr = SubElement(edge, 'data')
    magnitude_attr.set('key', 'magnitude')
    magnitude_attr.text = str(magnitude) if magnitude is not None else ''
    
    sign_attr = SubElement(edge, 'data')
    sign_attr.set('key', 'sign')
    sign_attr.text = str(sign) if sign is not None else ''
    
    p_value_attr = SubElement(edge, 'data')
    p_value_attr.set('key', 'p_value')
    p_value_attr.text = str(p_value) if p_value is not None else ''
    
    return edge


def main(input_file, output_file, p_value_threshold, attr_length, type_underscore):
    # Create an empty graph
    graph = Graph()

    with open(input_file, 'r') as f:
        next(f)  # Skip header row
        for line in f:
            if not (line.rstrip().endswith("\tNaN") or line.rstrip().endswith("\t0")):
                fields = line.strip().split('\t')

                if type_underscore:
                    node_type_1 = fields[0].split('_')[0]  # Extract everything before the first underscore as type for the first attribute
                    attribute_name_1 = '_'.join(fields[0].split('_')[1:])  # Extract everything after the first underscore as name for the first attribute

                    node_type_2 = fields[1].split('_')[0]  # Extract everything before the first underscore as type for the second attribute
                    attribute_name_2 = '_'.join(fields[1].split('_')[1:])  # Extract everything after the first underscore as name for the second attribute
                
                else:
                    node_type_1 = "Node"
                    node_type_2 = "Node"

                    attribute_name_1 = fields[0]
                    attribute_name_2 = fields[1]

                x2_value = fields[-2]
                magnitude = abs(float(fields[-2]))
                sign = '-' if float(fields[-2]) < 0 else '+'
                p_value = fields[-1]

                if float(p_value) <= p_value_threshold:
                    # Add nodes and edges to the graph
                    node1 = graph.add_node(f'{node_type_1}_{attribute_name_1}')
                    node2 = graph.add_node(f'{node_type_2}_{attribute_name_2}')
                    edge = graph.add_edge(node1, node2)

                    # Add attributes to the node
                    node1['type'] = node_type_1
                    node2['type'] = node_type_2
                    node1['name'] = attribute_name_1
                    node2['name'] = attribute_name_2
                    node1['truncated_name'] = (attribute_name_1[:attr_length] + '..') if len(attribute_name_1) > attr_length else attribute_name_1
                    node2['truncated_name'] = (attribute_name_2[:attr_length] + '..') if len(attribute_name_2) > attr_length else attribute_name_2

                    # Add attributes to the edge
                    edge['edge_type'] = 'Same_type' if node_type_1 == node_type_2 else 'Different_type'
                    edge['x2_value'] = x2_value
                    edge['magnitude'] = magnitude
                    edge['sign'] = sign
                    edge['p_value'] = p_value

    # Write the graph to a GraphML file
    GMLparser = GraphMLParser()
    GMLparser.write(graph, output_file)

if __name__ == '__main__':
    parser = argparse.ArgumentParser(description='Convert tab-separated file to GraphML format.')
    parser.add_argument('input_file', type=str, help='Path to the input tab-separated file.')
    parser.add_argument('output_file', type=str, help='Path to save the output GraphML file.')
    parser.add_argument('--p_value_threshold', type=float, default=0.005, help='P-value threshold for edge creation.')
    parser.add_argument('--attribute_name_length', type=int, default=10, help='Length to truncate attribute names.')
    parser.add_argument('--type_underscore', action='store_true', help='Assume attribute name before first underscore identifies its type.')

    args = parser.parse_args()

    # Since `type_underscore` is a boolean flag, it will be False if not present
    main(args.input_file, args.output_file, args.p_value_threshold, args.attribute_name_length, args.type_underscore)
```
