## Supplementary material for "ParallelEvolCCM: Quantifying co-evolutionary patterns among genomic features": Documentation, source code, and example files used in the manuscript.: PECCM_BuildFeatureHistogram.R

```
# Load necessary library
if (!require(ggplot2)) {
  install.packages("ggplot2")
  library(ggplot2)
}

# Read command line arguments
args <- commandArgs(trailingOnly = TRUE)

# Check for proper usage
if (length(args) != 1) {
  stop("Usage: Rscript script_name.R input_file.tsv")
}

# Get the input file path from the command line
input_file <- args[1]

# Read the tab-separated input file
feature_matrix <- read.table(input_file, header = TRUE, sep = "\t", row.names = 1)

# Check if data is correctly loaded and processed
if (nrow(feature_matrix) == 0 || ncol(feature_matrix) == 0) {
  stop("Error: The input file doesn't contain valid data.")
}

text_theme <- theme(
  plot.title = element_text(size = 20, face = "bold"),  # Title size
  axis.title = element_text(size = 30),                # Axis title size
  axis.text = element_text(size = 30)                  # Axis tick label size
)

# Compute the feature count across genomes
feature_counts <- colSums(feature_matrix)

histogram <- ggplot(data.frame(feature_counts), aes(x = feature_counts)) +
  geom_histogram(binwidth = 10, fill = "blue", color = "black") +
  labs(#title = "Histogram of Feature Counts Across Genomes",
       x = "Number of Genomes",
       y = "Frequency") +
  theme_minimal() + 
  theme(panel.background = element_rect(fill = "white", colour = NA), 
        text = element_text(size = 12, color = "black"),
        plot.background = element_rect(fill = "white", colour = NA),
        axis.text = element_text(color = "black"),
        panel.grid.major = element_line(color = "grey90"),
        panel.grid.minor = element_line(color = "grey95"),
        plot.margin = unit(c(0, 1, 0, 0), "cm")) +
  text_theme

        
# Save the histogram as a .png file
ggsave("a_xxx_feature_histogram.jpg", plot = histogram, width = 8, height = 6, units = "in")
```
