## Supplementary material for "ParallelEvolCCM: Quantifying co-evolutionary patterns among genomic features": Documentation, source code, and example files used in the manuscript.: PECCM_BuildStatHistogram.R

```
# Load necessary library
if (!require(ggplot2)) {
  install.packages("ggplot2")
  library(ggplot2)
}

# Read command line arguments
args <- commandArgs(trailingOnly = TRUE)

# Check for proper usage
if (length(args) != 1) {
  stop("Usage: Rscript script_name.R input_file.tsv")
}

# Get the input file path from the command line
input_file <- args[1]

# Read the tab-separated input file, skipping the header
data <- read.table(input_file, header = TRUE, sep = "\t")

# Extract the second-last (X2 statistics) and last (p-values) columns
x2_values <- data[[ncol(data) - 1]]
p_values <- data[[ncol(data)]]

# Filter out rows where p-value is zero or X2 or p-values are NA
valid_indices <- !is.na(x2_values) & !is.na(p_values) & p_values != 0
x2_values <- x2_values[valid_indices]
p_values <- p_values[valid_indices]

text_theme <- theme(
  plot.title = element_text(size = 20, face = "bold"),  # Title size
  axis.title = element_text(size = 30),                # Axis title size
  axis.text = element_text(size = 30)                  # Axis tick label size
)

# Create a histogram for X2 values
x2_hist <- ggplot(data.frame(X2 = x2_values), aes(x = X2)) +
  geom_histogram(binwidth = 1, fill = "blue", color = "black") +
  labs(
       x = "X² Value",
       y = "Frequency") +
  theme_minimal() +
  theme(panel.background = element_rect(fill = "white", color = NA),
        plot.background = element_rect(fill = "white", color = NA),
        axis.text = element_text(color = "black"),
        panel.border = element_blank(),               # Remove the panel border
        plot.margin = unit(c(0, 0, 0, 0), "cm")) +
  text_theme

# Save the X2 histogram as a JPG file
ggsave("b_xxx_x2_histogram.jpg", plot = x2_hist, width = 8, height = 6, units = "in")

# Create a histogram for p-values
p_hist <- ggplot(data.frame(p = p_values), aes(x = p)) +
  geom_histogram(binwidth = 0.05, fill = "red", color = "black") +
  labs(
       x = "p-value",
       y = "Frequency") +
  theme_minimal() +
  theme(panel.background = element_rect(fill = "white", color = NA),
        plot.background = element_rect(fill = "white", color = NA),
        axis.text = element_text(color = "black"),
        panel.border = element_blank(),               # Remove the panel border
        plot.margin = unit(c(0, 0, 0, 0), "cm")) +
  text_theme

# Save the p-value histogram as a JPG file
ggsave("c_xxx_p_histogram.jpg", plot = p_hist, width = 8, height = 6, units = "in")

# Filter p-values to include only those <= 0.1
filtered_p_values <- p_values[p_values <= 0.1]

# Create a histogram for p-values <= 0.1, using log scale for the x-axis
log_x_hist <- ggplot(data.frame(p = filtered_p_values), aes(x = p)) +
  geom_histogram(binwidth = 0.1, fill = "green", color = "black") +
  scale_x_log10() +
  labs(
       x = "p-value",
       y = "Frequency") +
  theme_minimal() +
  theme(panel.background = element_rect(fill = "white", color = NA),
        plot.background = element_rect(fill = "white", color = NA),
        axis.text = element_text(color = "black"),
        panel.border = element_blank(),               # Remove the panel border
        plot.margin = unit(c(0, 0, 0, 0), "cm")) +
  text_theme

# Save the log-scaled p-value histogram as a JPG file
ggsave("d_xxx_filtered_p_histogram.jpg", plot = log_x_hist, width = 8, height = 6, units = "in")
```
