## Supplementary figures and images for "ParallelEvolCCM: Quantifying co-evolutionary patterns among genomic features"

### a_1000_feature_histogram.jpg

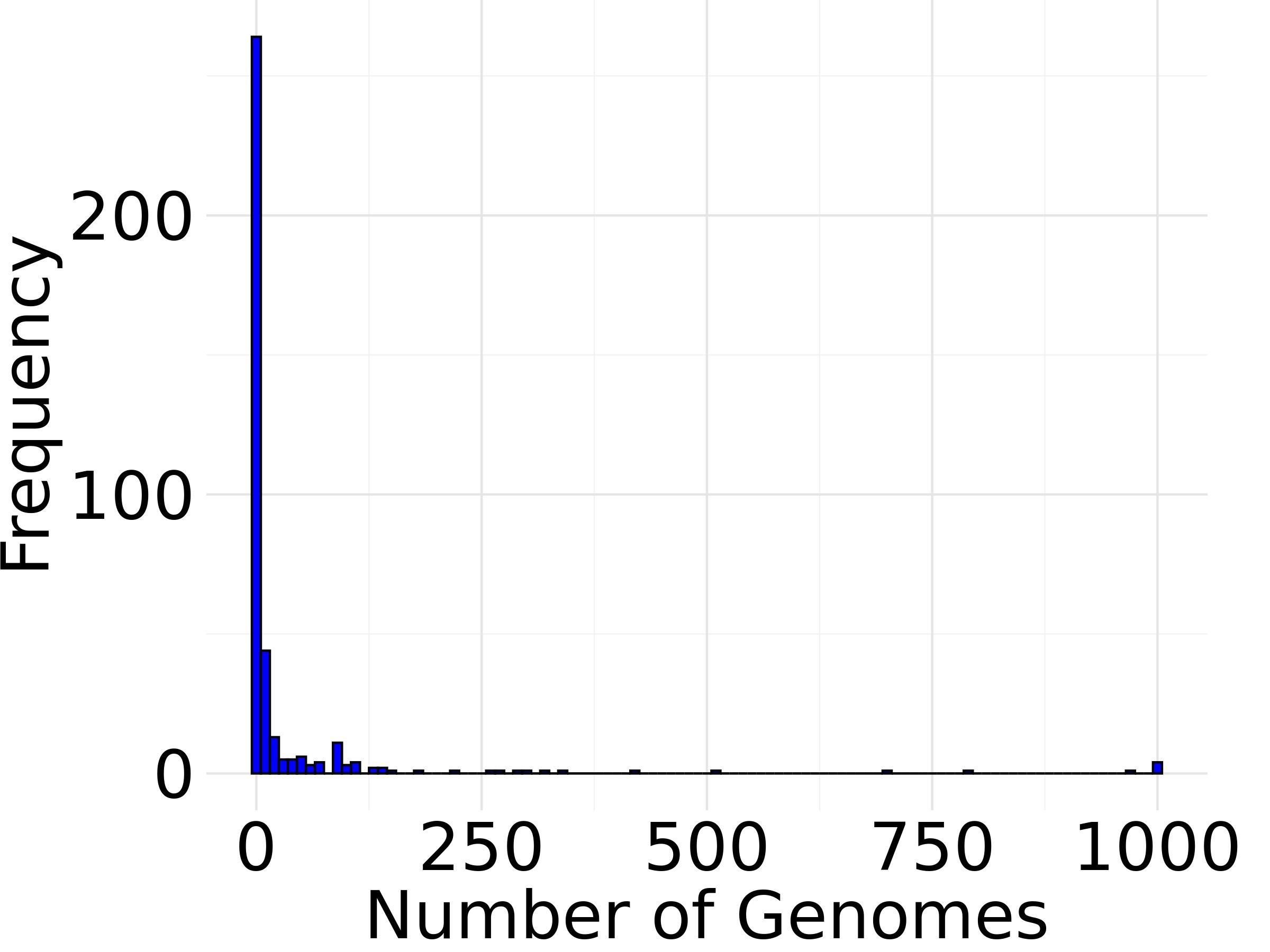

### a_xxx_feature_histogram.jpg

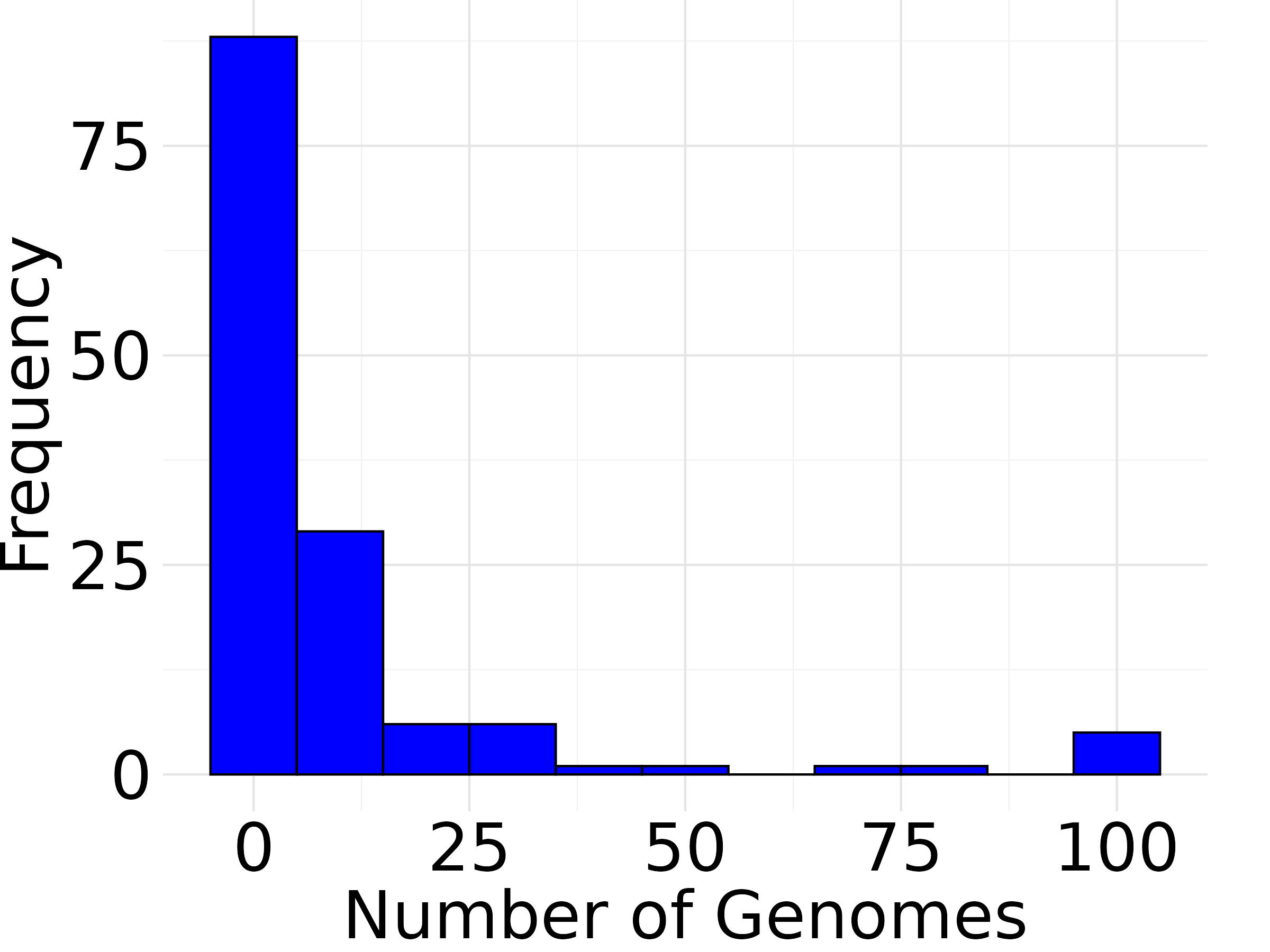

### b_1000_x2_histogram.jpg

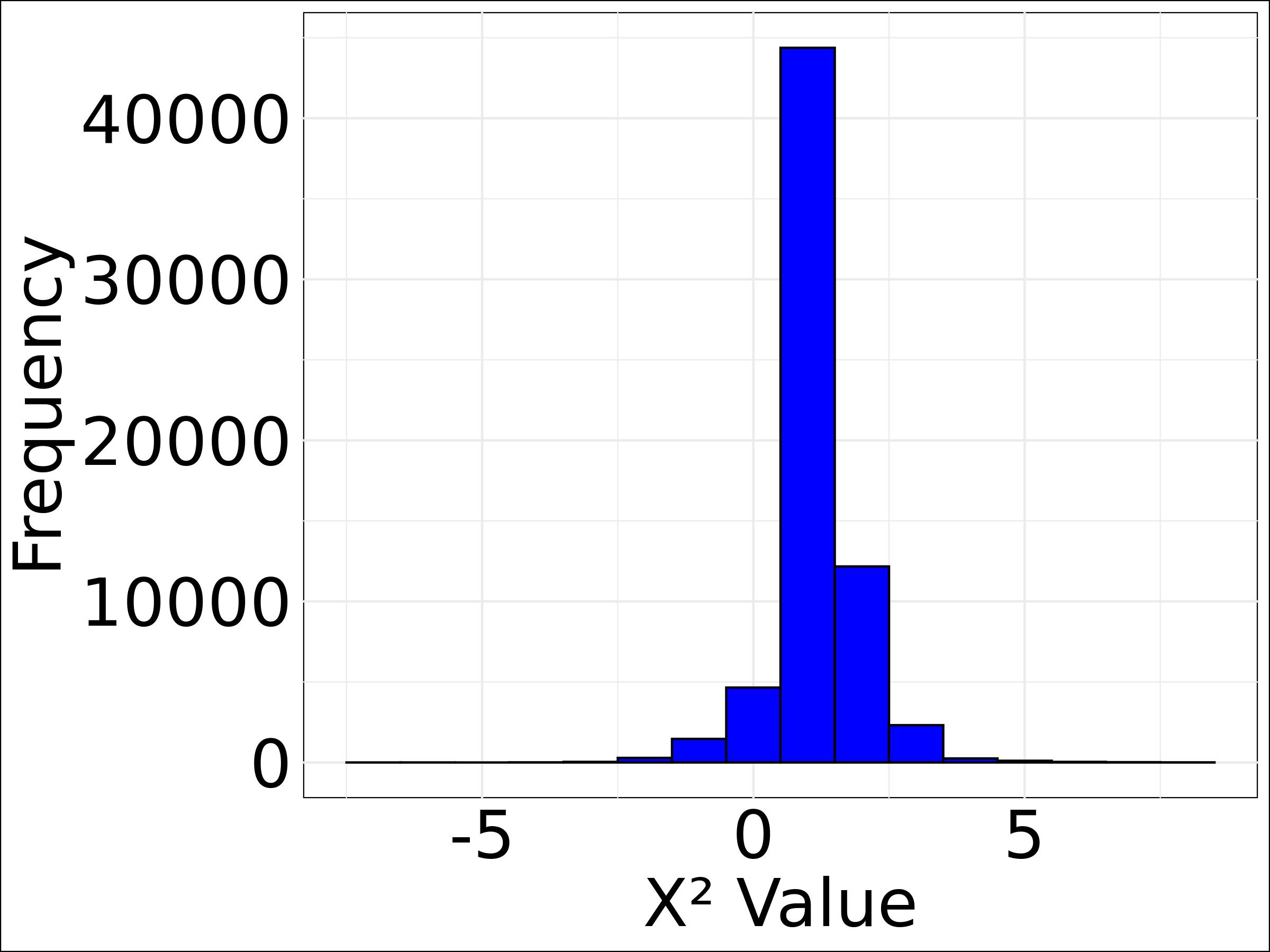

### b_xxx_x2_histogram.jpg

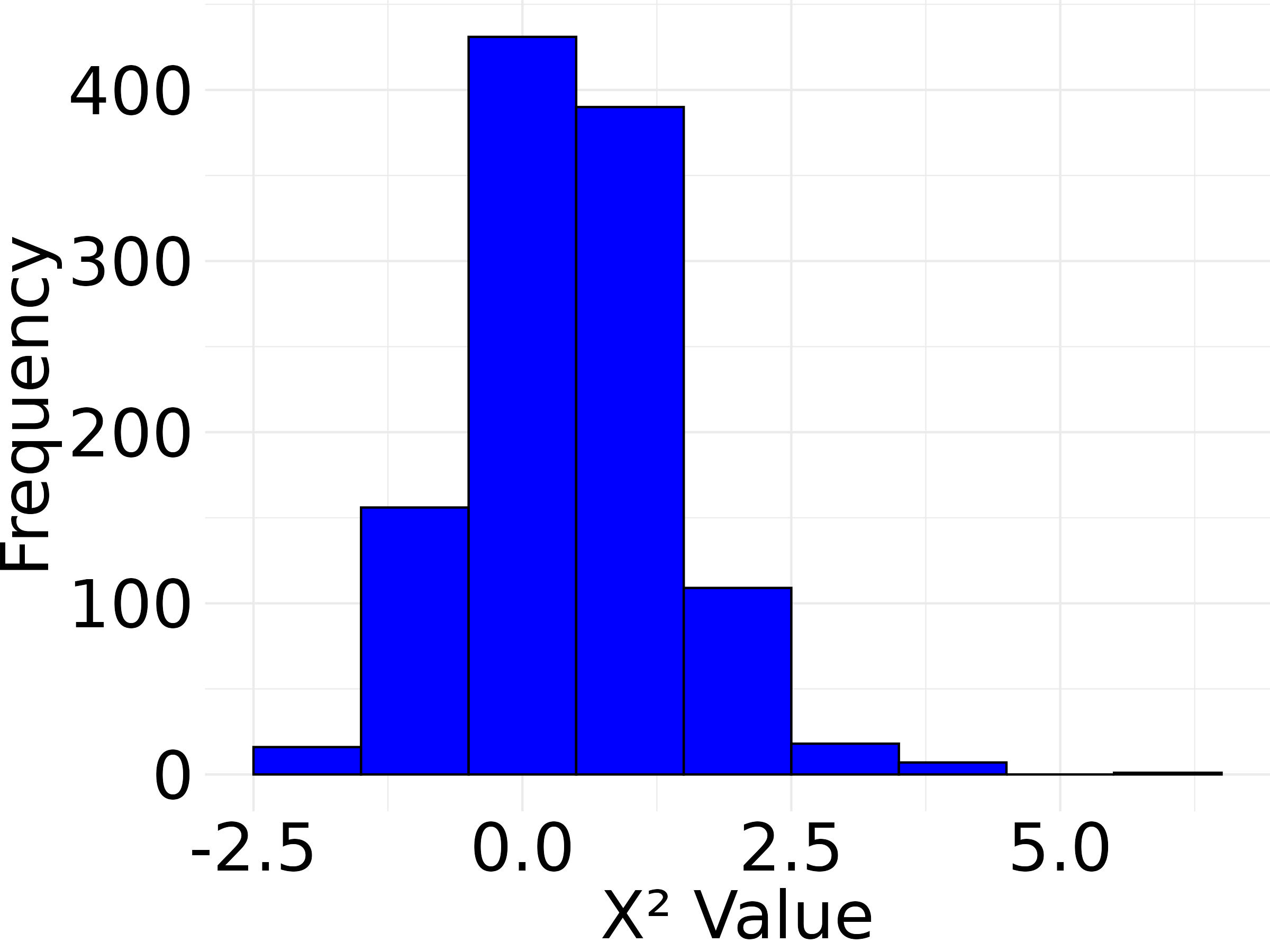

### c_1000_p_histogram.jpg

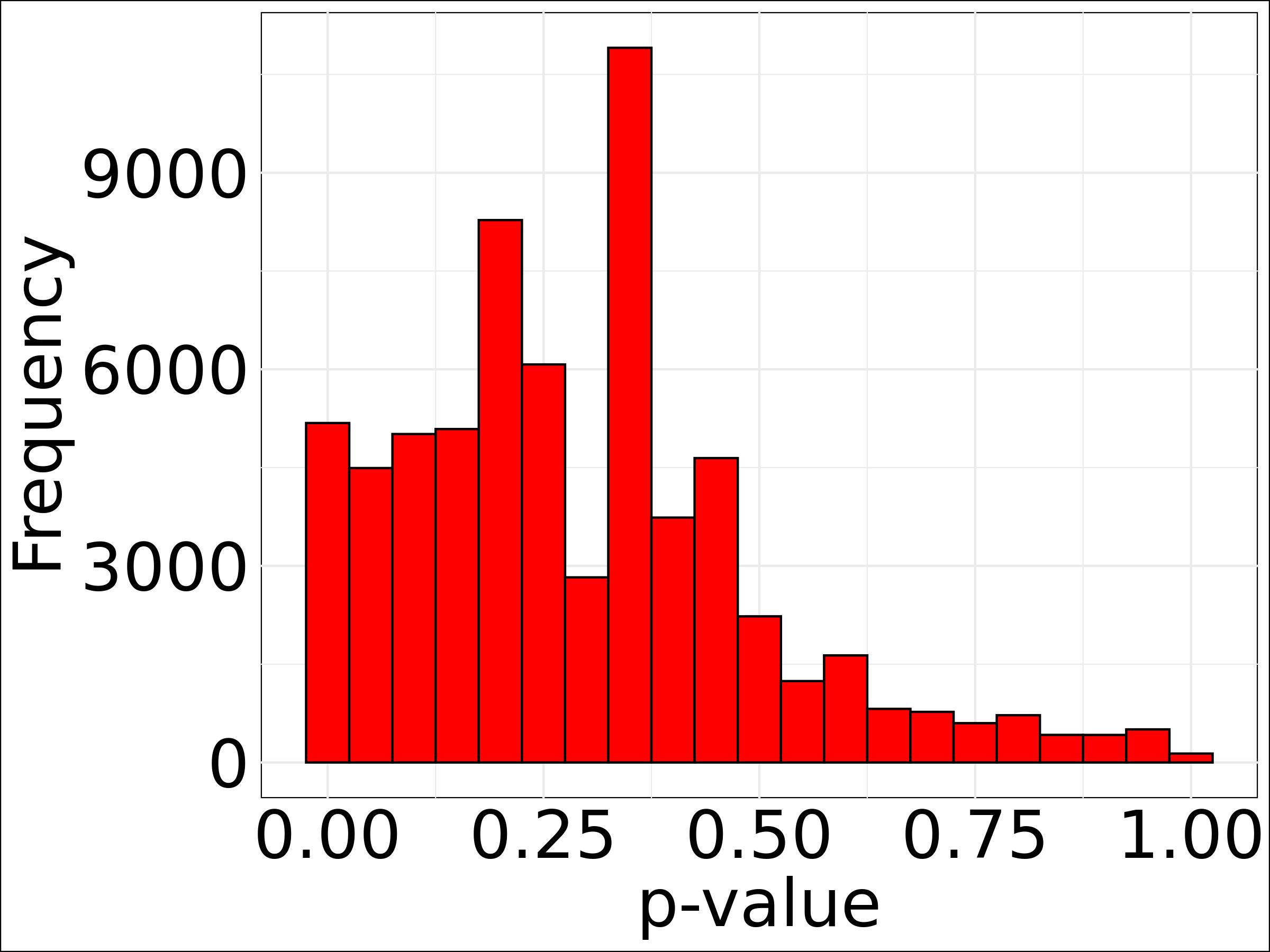

### c_xxx_p_histogram.jpg

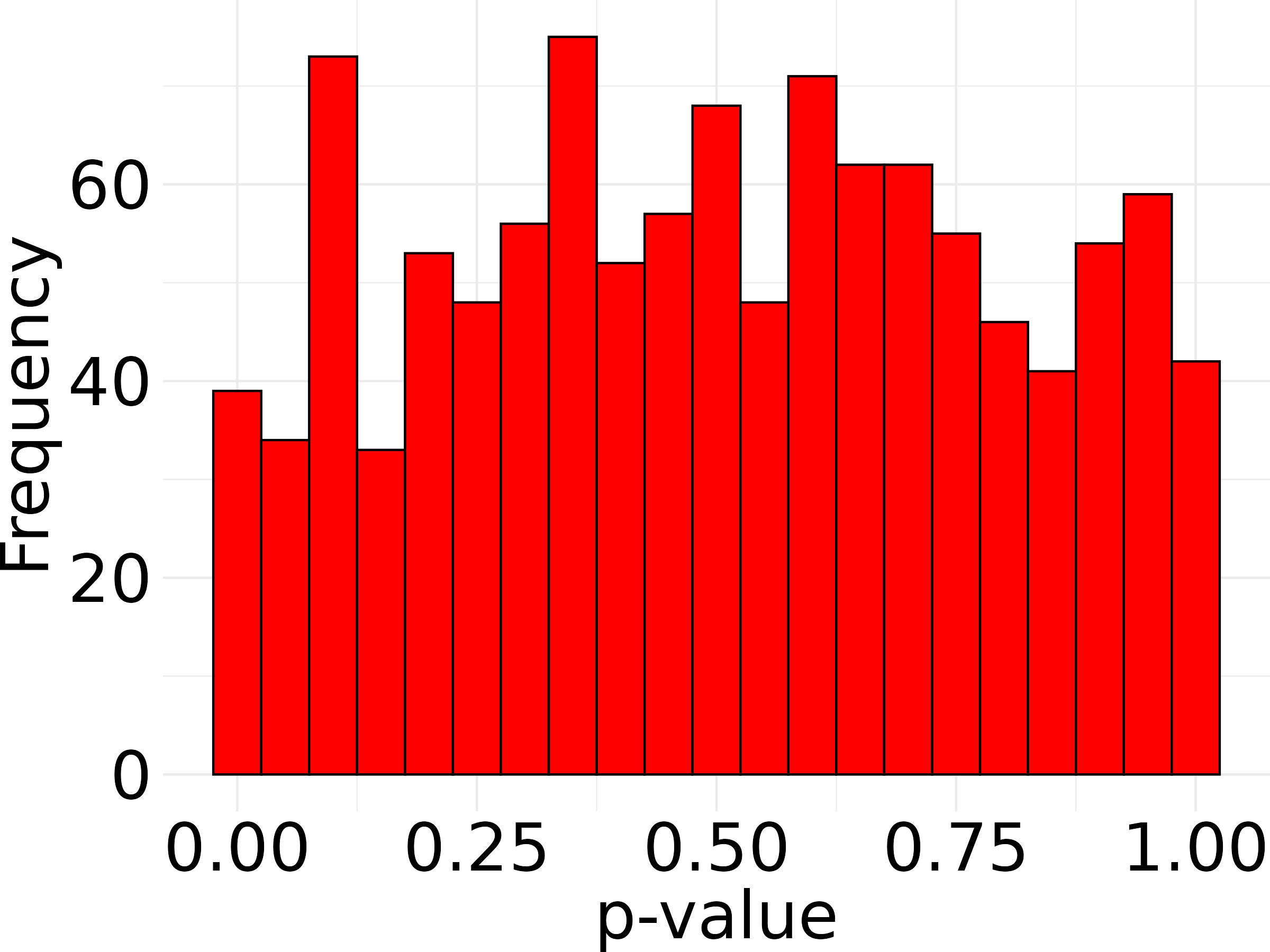

### d_1000_filtered_p_histogram.jpg

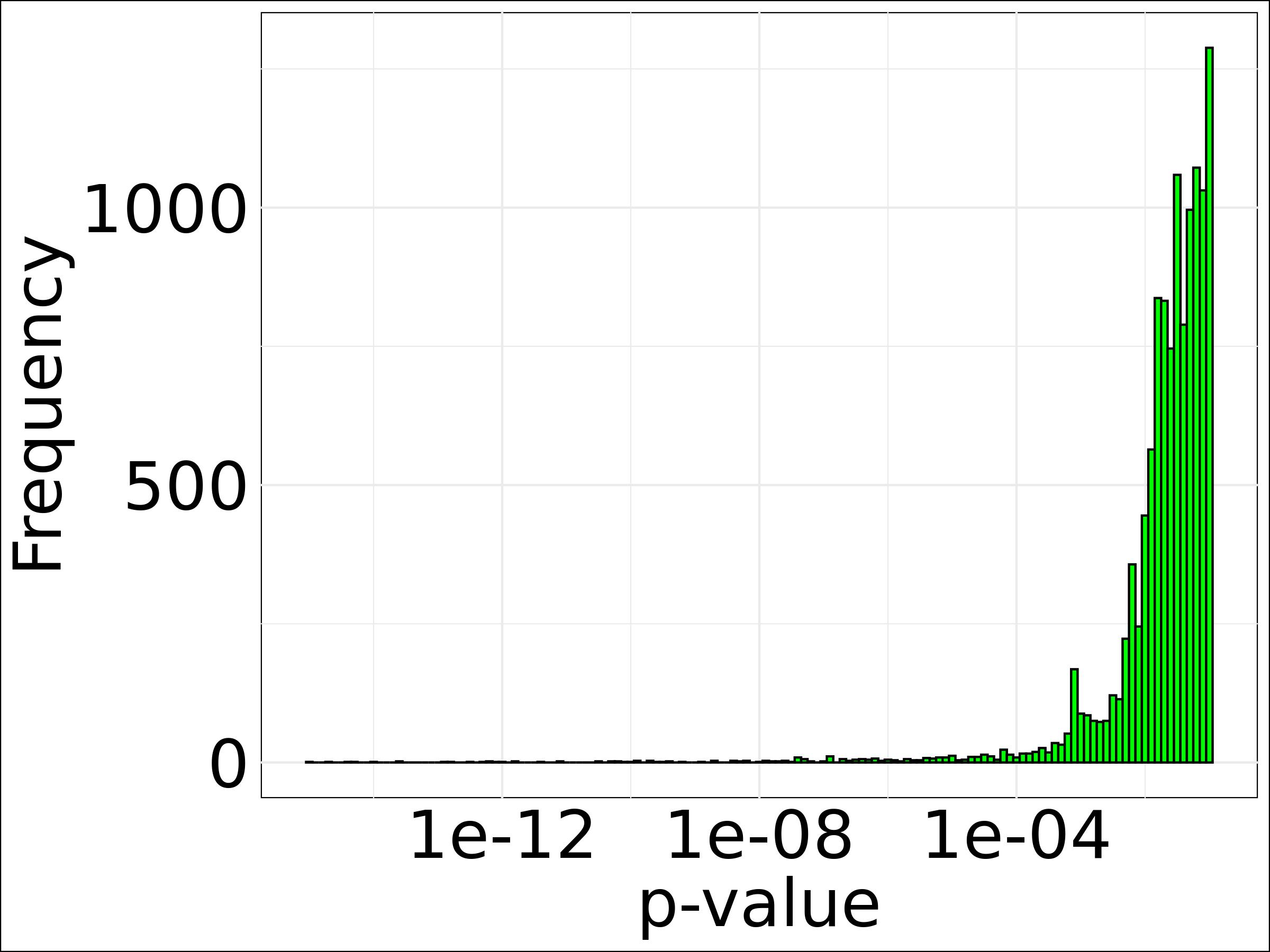

### d_xxx_filtered_p_histogram.jpg

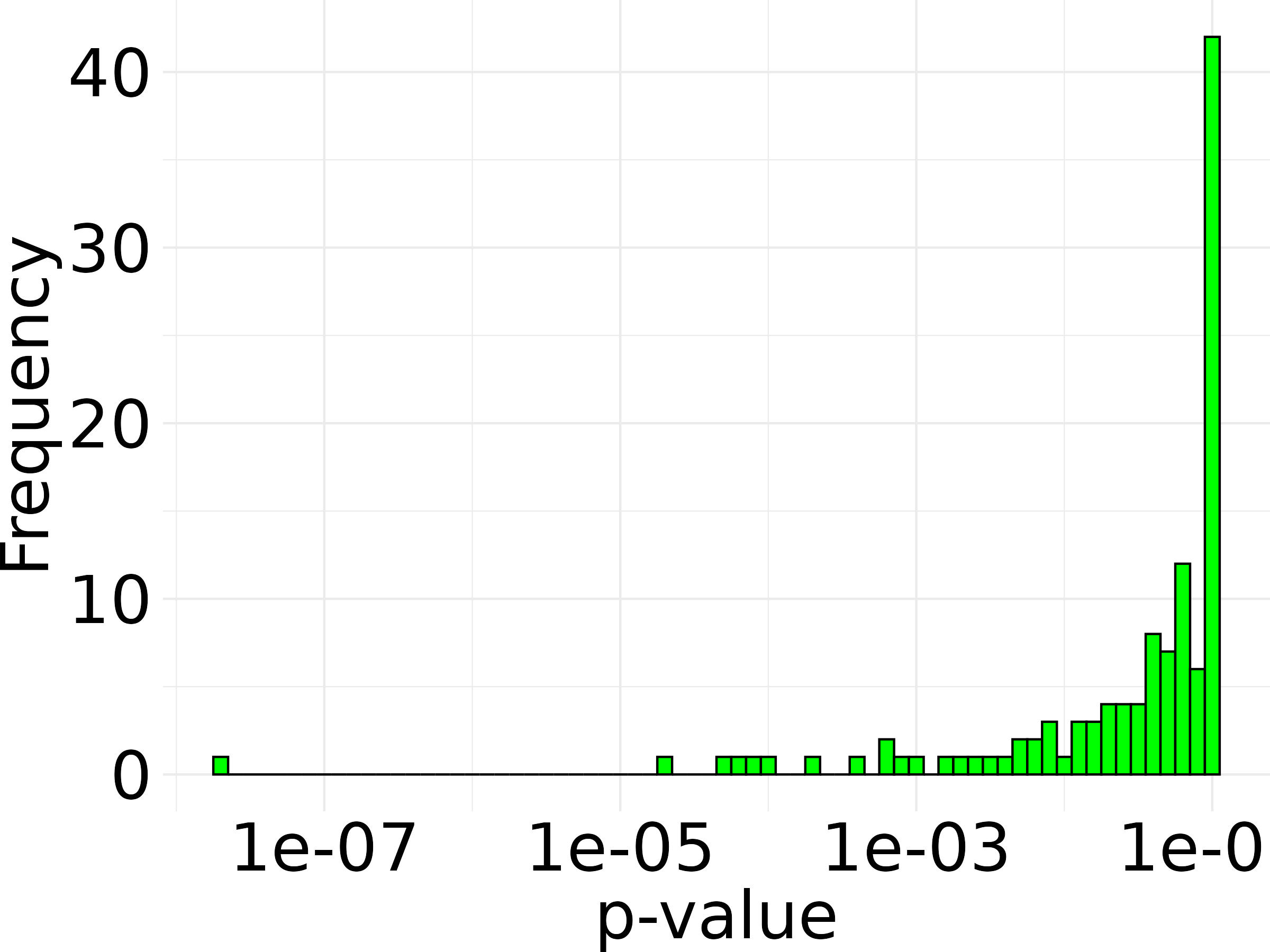
